# Enhancing sleep via rocking ameliorates motor behavior and reduces beta-amyloid levels in a mouse model of Alzheimer’s disease

**DOI:** 10.1101/2024.07.15.603512

**Authors:** Luyan Zhang, Letizia Santoni, Nam Anh Ngo, Reyila Simayi, Eleonora Ficiará, Luisa de Vivo, Michele Bellesi

## Abstract

Alzheimer’s disease (AD) is a progressive neurodegenerative disorder associated with cognitive decline and characterized by beta-amyloid plaque and tau tangle pathology. Recent research indicates a bidirectional relationship between AD pathology and sleep disturbances, with disrupted sleep exacerbating AD progression through increased beta-amyloid and tau accumulation. This strongly indicates that improving sleep may exert a direct protective effect on preventing the accumulation and spreading of AD pathology, and possibly slow the cognitive decline.

Here we investigated the effects of enhancing sleep via vestibular stimulation (rocking) on AD progression in a 3xTg mouse model. Over a four-month period starting in early adulthood (p60), we monitored sleep patterns, motor function, memory, and AD pathology. Twelve-hour rocking during the light period significantly increased non-rapid eye movement (NREM) sleep duration in mice, although this effect diminished over time due to habituation. Despite this, rocking attenuated motor function decline and reduced beta-amyloid levels in the cerebral cortex of treated mice. No noticeable changes in tau levels were observed following sleep enhancement.

In conclusion, our findings highlight the potential of non-pharmacological methods to enhance NREM sleep and modify disease trajectory in AD models, emphasizing the critical role of sleep in neuroprotection.

## INTRODUCTION

Alzheimer’s disease (AD) is a progressive neurological condition and the primary cause of dementia [1]. Recent studies suggest a two-way relationship between AD pathology and sleep disturbances [2]. Individuals with AD commonly experience disrupted sleep, marked by frequent nighttime awakenings and difficulty maintaining sleep. Conversely, chronically disturbed sleep raises the risk of AD and leads to significant increases in beta-amyloid and hyperphosphorylated tau levels, also driving tau pathology spreading in the brains of mice and humans [3–6]. Moreover, the specific reduction in NREM sleep has been associated with tau pathology in people with normal cognition or very mild cognitive impairment [7,8].

While the exact mechanism remains unclear, emerging research indicates that disrupted sleep compromises the glymphatic system’s function—a glial-dependent waste clearance pathway in the brain [9,10]. This system clears soluble waste proteins and metabolic byproducts, including beta-amyloid and tau, preferentially during sleep. Consequently, insufficient NREM sleep may impair the glymphatic system function, thus limiting the removal of extracellular beta-amyloid and tau and favoring their accumulation [9]. Conversely, NREM sleep could exert a direct protective effect on preventing the accumulation and spreading of pathological amyloid and tau. This emphasizes the potential of enhancing NREM sleep to mitigate AD progression.

Enhancing NREM sleep is commonly attempted through pharmaceutical interventions [11]. However, drugs targeting GABAergic neurotransmission, such as gamma-hydroxybutyrate (GHB), gaboxadol, and tiagabine, though effective in increasing NREM sleep, pose risks of drug dependence, tolerance, and daytime side effects like somnolence. Moreover, they often fail to yield corresponding memory benefits in older adults and may even have amnestic effects [12–15]. Alternatively, non-pharmacological methods have been explored, including magnetic or weak electric currents applied to the scalp during sleep, which can boost sleep slow waves [16,17]. However, these methods, while effective, may be impractical or potentially hazardous with repeated application. Consequently, recent research has shifted focus towards more natural means of enhancing NREM sleep, such as sensory stimulation techniques [11]. Vestibular stimulation was among the first techniques explored for its potential to induce sleep, likely due to the traditional belief that physical rocking aids in sleep induction, as seen in babies or hammock swinging. Research findings from studies in both humans and rodents have demonstrated that rocking not only facilitates sleep onset but also increases the duration of NREM sleep without impacting REM sleep [18–20].

In this study, we investigated whether enhancing sleep through rocking could delay the progression of AD in a mouse model harboring familial AD mutations, characterized by both plaque and tangle pathology. We assessed the effects of rocking on sleep patterns, motor function, memory, and AD pathology over a four-month treatment period, starting in early adulthood.

## MATERIALS AND METHODS

### Animals

Five-week-old B6;129-Tg (APPSwe, tauP301L)1Lfa Psen1 tm1Mpm/Mmjax (3xTg, weight 25–35 g) mice were acquired from the Jackson Laboratory (Bar Harbor, ME USA 04609). All mice were housed in groups of four in environmentally controlled cages and subsequently placed in sleep recording chambers. Mice were under 12 h dark/light cycle with lights on at 8:00 P.M., at the temperature of 24 ± 1 °C, with food and water available ad libitum and replaced daily at 8:00 A.M. All procedures were in accordance with the guidelines laid down by the European Communities Council Directives (2010/63/EU) for the care and use of laboratory animals under an approved protocol (609/2023-PR) by Veterinary Health Dept. of the Italian Ministry of Health. All appropriate measures were taken to minimize pain and discomfort in experimental animals.

### Experimental Design

Male mice were divided into two weight-balanced groups, the sleep enhancement (SE) and normal sleep (S) group. SE was achieved by rocking the animals on a shaker that allowed horizontal movements at 1Hz, for 12 hours/day during the light period. This procedure has been shown to increase the time spent in NREM sleep at the expenses of wake time. Mice in the S group were positioned within the identical sleep apparatus used for the SE mice. However, they remained undisturbed with the rocking system turned off. Rocking started following a 10-day acclimatization period, with the final three days serving as the baseline. In a pilot set of experiments aimed at determining the optimal duration of the chronic stimulation, SE mice were rocked during the light period for 12 hours for 20 consecutive days. In the main experiment, we opted for a different paradigm of stimulation by introducing a block design (one week of stimulation always during the light period only followed by one week of no stimulation for nine cycles – 18 weeks overall, starting at P60). Throughout the whole experiment, all the mice were video monitored for sleep detection using motion activity and were behaviorally evaluated biweekly using hindlimb clasping test and ledge walking test.

### Electroencephalographic recordings

A subset of mice was implanted with electrodes for electroencephalographic recordings and was used to test the efficacy of the rocking procedure in this mouse model. Briefly, mice were anesthetized with isoflurane (2.5% for induction and 1.5% for maintenance) and implanted with frontal and parietal electrodes to monitor brain activity. An additional electrode was placed above the cerebellum and used as reference. To measure muscle activity, a pair of silver wires were placed into the muscles on both sides of the back of the neck. Implanted mice were kept individually in transparent Plexiglas cages throughout the experiment. After recovery from the surgery, recordings started at the beginning of the light phase. Data acquisition was carried out continuously with an Open Ephys system across the 24h per 2 days. The signal was filtered at 0.1-40 Hz for EEG and 10-50 Hz for EMG. The paradigm of acquisition consisted in 1 day of baseline and one day of sleep enhancement via rocking at 1Hz during the light phase. EEG scoring was performed offline by visual inspection of 4s epochs using Sleep Sign software. Behavioral state and power analysis were analyzed using custom-made Matlab scripts. Slow wave activity (SWA) was computed by averaging the EEG NREM sleep power spectrum between 0.5 and 4Hz. Cumulative SWA (slow wave energy, [SWE]) was calculated by cumulative summing NREM sleep SWA over 12h of the light period. Sleep episodes were defined as periods of NREM sleep longer than 8 seconds. Short periods of wake shorter than 16 seconds in between NREM sleep episodes of at least 20 seconds were identified as brief arousals.

### Sleep quantification using motion detection

To avoid possible tissue damage and inflammation resulting from the implant of EEG electrodes, in the pilot and main experiment sleeping and waking states were inferred by the analysis of motion activity obtained by continuous video monitoring with infrared cameras. As described previously [21,22], this method consistently estimates total sleep time with ≥90% accuracy even though it cannot distinguish NREM sleep from REM sleep.

### Ledge Walking and Hindlimb Clasping Test

The ledge walking was used to test motor coordination, which is usually impaired in mice models of neurodegenerative disorders. The test was performed by placing a mouse on the cage’s ledge. Mice typically walk along the ledge and attempt to descend back into the cage. We assigned a test score (0 to 3) based on walking posture. Normal (score 0); slight imbalance while walking (score 1); ineffectively use of hind legs or landing on the head (score 2); falling off or extreme reluctance to move (score 3). Hindlimb clasping is a simple test that assesses hindlimb position relative to the body and is used to evaluate disease progression in mouse models of neurodegenerative diseases [23,24]. The mice were lifted by grasping the tail near its base and observed for 10 seconds. A score (0 to 3) was gives as it follows: hindlimb consistently splayed outward (score 0), one hindlimb retracted >5 s of the time (score 1), both hindlimbs retracted >5 s (score 2), both hindlimbs fully retracted and touch the abdomen >5 s (score 3). Each of these tests was conducted three times, and the average score was calculated for each time point.

### Novel Object Recognition

After the 18 weeks of sleep manipulation, all mice were tested with a novel-object recognition (NOR) test. NOR was performed in an open field with a camera mounted above the open field recorded the movements of the mouse throughout the trial. The NOR test had three phases: the habituation phase, where mice freely investigate the box for 5 minutes; the training phase, involving a 10-minute exposure to a box with two identical objects; and the testing phase, occurring after a 3-hour intertrial pause, where mice explore a box containing one novel and one known object for 10 minutes. The box was cleaned using 75% ethanol between each trial. The time spent exploring each object a discrimination score was calculated with the following formula: discrimination score = time exploring the novel object − time exploring the familiar object) / (total time exploring both objects.

### Tissue Collection

The next day following the NOR test, all mice were sacrificed between 9:00 and 11:00 A.M. to maintain the time of tissue collection within the same 2-h time of day window for all experimental groups. Mice were briefly anesthetized with isoflurane (1–1.5% volume) and sacrificed by cervical dislocation. Brains were extracted, quickly dissected to separate the cortex, were immediately frozen in dry ice, and were used for molecular evaluations. Mice used for immunohistochemistry were perfused intracardiacally with a solution containing phosphate buffer and paraformaldehyde 4%. Brains were post-fixed for 24 hours in the same fixative and then sliced in 50 µm coronal sections using a vibratome.

### Western Blotting

Through a glass/glass tissue homogenizer, mouse cortex tissue samples were homogenized in freezing homogenization buffer containing 10 mM HEPES, 1.0 mM EDTA (Sigma), 2.0 mM EGTA (Promega), 0.5 mM DTT (Boston BioProducts), 0.1 mM PMSF (Invitrogen), Protease Inhibitor Cocktail (Fluka), 100 nM microcystin (Roche). Next, 500 μL of each sample was boiled in 50 μL of Sodium Dodecyl Sulfate (SDS) at 90/95 °C for 8 min and kept at – 80 °C. The protein concentration was evaluated with bicinchoninic acid assay. One sample was discarded for technical problems during the preparation.

Equal amounts of protein from each animal (15μg for beta-amyloid, 5μg for total tau, and 14 μg for p-tau) were loaded onto the same gels with sample loading order randomized. Homogenate samples of each S and SE mice were resolved by Tris-HCl gel electrophoresis in 1X Tris/Glycine/SDS running buffer. Then, the proteins were transferred onto 0.45 μm pore size nitrocellulose membranes in 1X Tris/Glycine/Methanol transfer buffer. After transfer, membranes were stained with Ponceau S, acquired for total protein quantification at the Chemidoc (Bio-Rad), and then washed 3 times with phosphate buffer solution (PBS) + 0.1% Tween-20. The membranes were immunoblotted as follows. First, they were blocked in 5% non-fat dry milk in PBS + 0.1% Tween-20 with gentle shaking for 1h at room temperature; then, they were incubated for 2 h at room temperature with gentle shaking and overnight at 4 °C with one of the following primary antibodies: anti-beta-amyloid 1-42 antibody (beta-amyloid, RRID:AB_91677 Millipore #AB5078P, 1:1000), anti-tau antibody, mouse monoclonal (TAU, RRID:AB_477595 Sigma #T9450, 1:2000) or anti-pTau antibody (AT8 RRID:AB_223647 Thermo Fisher Scientific MN1020 or pTau-s404 Abcam #ab196364) diluted in PBS + 0.1% Tween-20 with 5% bovine serum albumin. Next, membranes were rinsed 3 times in PBS + 0.1% Tween-20 and then incubated with one of the following secondary antibodies: goat anti-rabbit IgG (1:10000) or Anti-mouse IgG (1:5000), HRP-linked antibody diluted in PBS + 0.1% Tween-20 with 3% non-fat dry milk for 90 min at room temperature with mild oscillation. Membranes were washed 3 times in PBS + 0.1% Tween-20 and, in the end, with ddH2O, incubated with enhanced ECL Chemiluminescence Reagent (ECL-Prime, GE Healthcare), and the bands were revealed by exposition to Molecular Imager Chemidoc XRS+ (Bio-Rad). The optical density of the bands was quantified by the ImageLab software (Bio-Rad). Since housekeeping proteins (e.g., β-actin and β-tubulin) can be affected by sleep and wake, they were not used as internal standard. Instead, optical density values were normalized to total protein loading obtained by the ponceau S staining [25] (normalized experimental signal = observed experimental signal/lane normalization factor; the lane normalization factor = observed signal of total protein for each lane/by the highest observed signal of total protein on the blot).

### Methoxy-X04 staining

Sections from both experimental conditions were washed with phosphate buffer and then exposed to Methoxy-X04 (1µg/ml, HB5252, Biozol) for 30 minutes to highlight beta-amyloid plaques. After staining sections were washed again, mounted on microscopy slides, and imaged using a Nikon confocal microscopy. Multiple random fields of cerebral cortex and hippocampus were analyzed for both groups of mice.

### Statistical Analysis

Statistical analysis was performed using GraphPad Prism software (La Jolla, CA, USA) and Matlab (The MathWorks Inc., Natick, MA, USA). Parametric statistics were used, alpha was set to 0.05 and appropriately corrected for multiple comparisons when required.

## RESULTS

### Rocking increases sleep duration, but its effect wanes over time

Previous research on C57B6 wild type mice demonstrated that rocking at a 1Hz frequency during the light period extended NREM sleep duration at the expense of wakefulness [19]. In an initial set of experiments, we aimed to verify the effectiveness of rocking in our 3xTg mice. To achieve this, we implanted EEG electrodes in six mice and recorded their EEG activity during a baseline period (no rocking) and an experimental day where they were rocked for 12 hours during the light period. Analysis of the sleep architecture showed that rocking increased time spent in NREM sleep and reduced wake time, while having no effect on REM sleep. This effect was limited to the light period when rocking was active (NREM: P<0.0001; REM: P=0.99; Wake: P<0.0001, Figure 1A-B), as behavioral states during the dark period remained unaffected (NREM: P=0.98; REM: P=0.99; Wake: P=0.92, Figure 1C). Further analysis of NREM episodes duration revealed that rocking predominantly increased the number of short to medium duration episodes (cluster 0-20 sec: P=0.0597; cluster 20-40 sec: P<0.0001, Figure 1D), indicating that rocking enhanced the propensity to fall asleep rather than extending the duration of long consolidate sleep episodes. Analysis of NREM sleep time course demonstrated that the effect of rocking was more pronounced at the beginning of light period rather than the end (bin 0-2 h: P<0.0001; bin 2-4: P=0.078, Figure 1E). The increase in NREM sleep did not significantly influence averaged SWA (2-way ANOVA 24h: F (1, 60) = 1.474e-013 P>0.99; 2-way ANOVA only light period: F (1, 30) = 2.751 P=0.11, Figure 1F), but it increased overall cumulative SWA over the 12h of rocking (SWE, P=0.011, Figure 1G). Finally, rocking did not increase neither the number of brief arousals (P=0.53, Figure 1H) nor the NREM sleep beta power (15 to 30Hz, P=0.19, Figure I).

**Figure 1.**
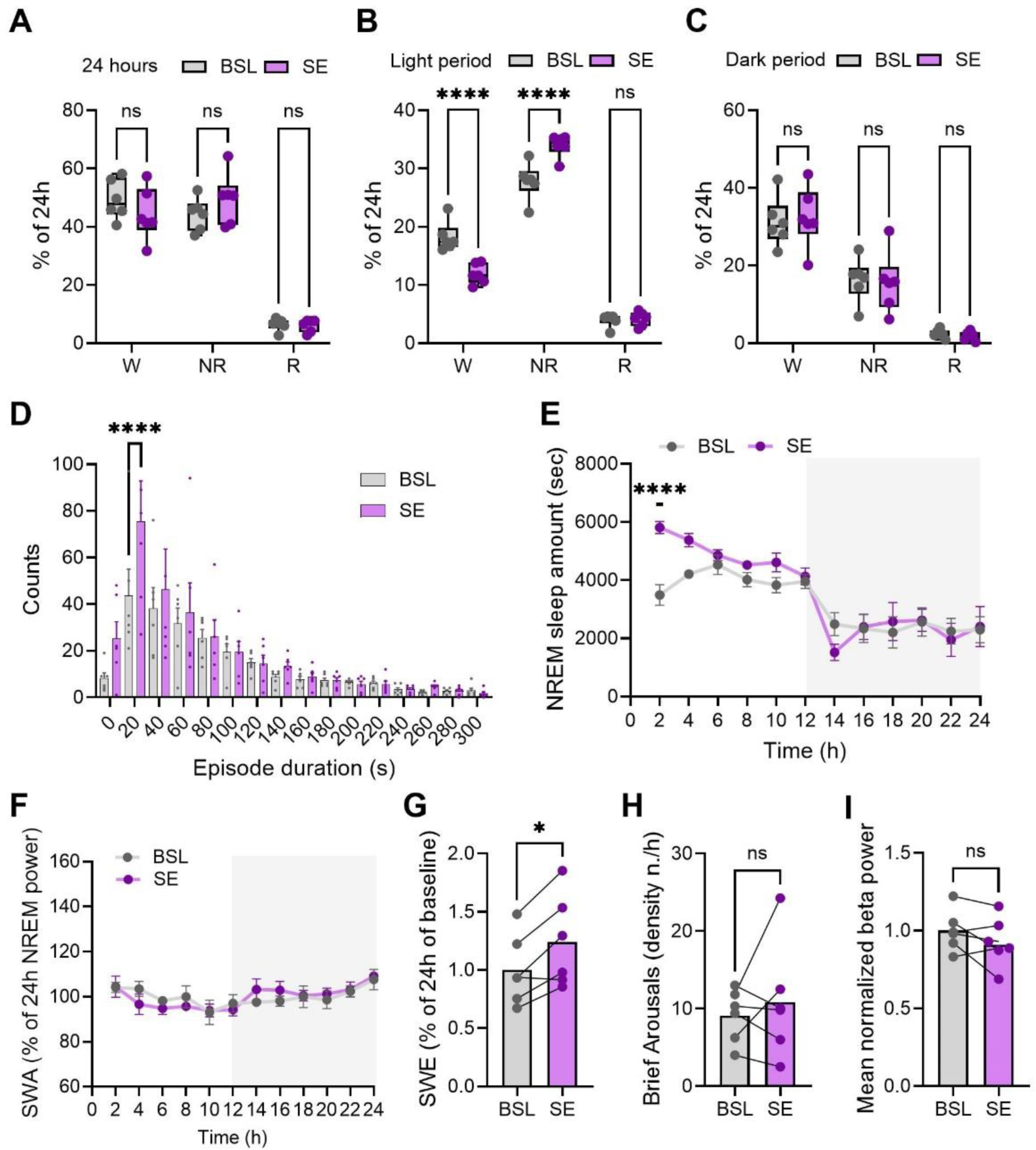
**A**. Distribution of wake (W), NREM sleep (NR), and REM sleep (R) amount in the 24h baseline (BSL, n=6) and sleep enhancement (SE, n=6). Dots represent individual mice. Ns: not significant. **B-C**. Distribution of W, NR, and R amount in the light (**B**) and dark (**C**) period of BSL and SE. Dots represent individual mice. ****P<0.0001. **D.** Frequency distribution of NREM sleep episodes for BSL and SE. Bars indicate mean ± SEM and dots represent individual mice. ****P<0.0001. **E.** Twenty-four hour NREM time course for BSL and SE. Values are mean ± SEM. Grey area indicates the dark period. ****P<0.0001. **F.** Twenty-four hour time course of NREM SWA for BSL and SE. Values are mean ± SEM. Grey area indicates the dark period. **G.** Cumulative SWA (SWE) over the 12h light cycle in BSL and SE. Bars indicate mean, and dots represent individual mice. *P<0.05. **H.** Density of brief arousals over the 12h light cycle for BSL and SE. Bars indicate mean, and dots represent individual mice. **I.** Normalized beta (15-30Hz) power over the 12h light cycle. Bars indicate mean, and dots represent individual mice.

In chronic experiments, we utilized sleep and wake quantification based on motion detection analysis to avoid keeping the mice implanted and tethered for extended periods, thereby reducing animal stress and the risk of inflammation and tissue reactions associated with electrode implants. We initially carried out a pilot experiment to establish the duration of the stimulation. In this experiment mice were rocked during for 12 hours during the light period for 20 consecutive days starting at post-natal day (p)40 (Figure 2A). Exposure to rocking increased the amount of sleep time in the first days, but this effect progressively faded off over the course of the stimulation, likely due to habituation (2-way ANOVA: F (1, 280) = 17.66 P<0.0001; Figure 2B). Sleep amount during the dark period did not differ between the two groups (2-way ANOVA: F (1, 280) = 2.547 P=0.1116; Figure 2C).

**Figure 2.**
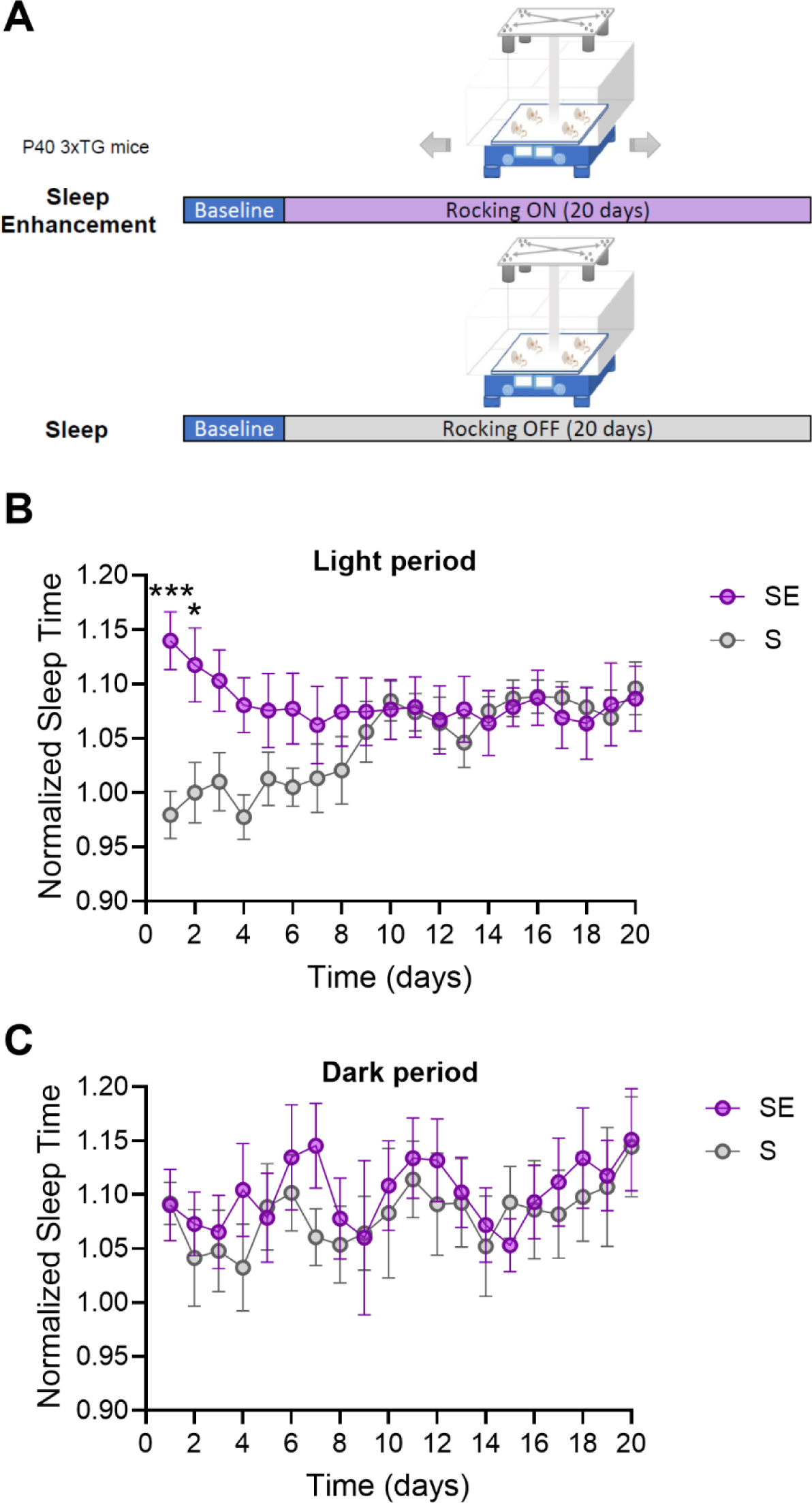
**A**. Experimental design of the pilot study. **B-C**. Sleep amount over the 20 days of observation period for light (**B**) and dark (**C**) period. Sleep amount is normalized to baseline mean sleep duration. (S, n=8; SE, n=8). Values are mean ± std. * P<0.05; ***P<0.001 (Sidak’s correction).

In following experiments, we modified the stimulation paradigm by implementing a block design, alternating between a week of stimulation and a week of no stimulation to contrast habituation. This pattern was repeated for 9 cycles over a total of 18 weeks, during which we continuously monitored sleep and wake via motion detection (Figure 3A). Analysis of sleep duration during the stimulation weeks for the SE group revealed an increase compared to baseline, while sleep duration remained stable in the control S group (Figure 3B). The effect of rocking was more pronounced initially and diminished over the course of the stimulation week (Day1 vs bsl: P=0.0069; Day2 vs bsl: P=0.019; Figure 3C). Notably, rocking did not affect sleep duration during the dark phase (RM-1way ANOVA: F (1.989, 13.92) = 0.95 P=0.4096; Figure 3D).

**Figure 3.**
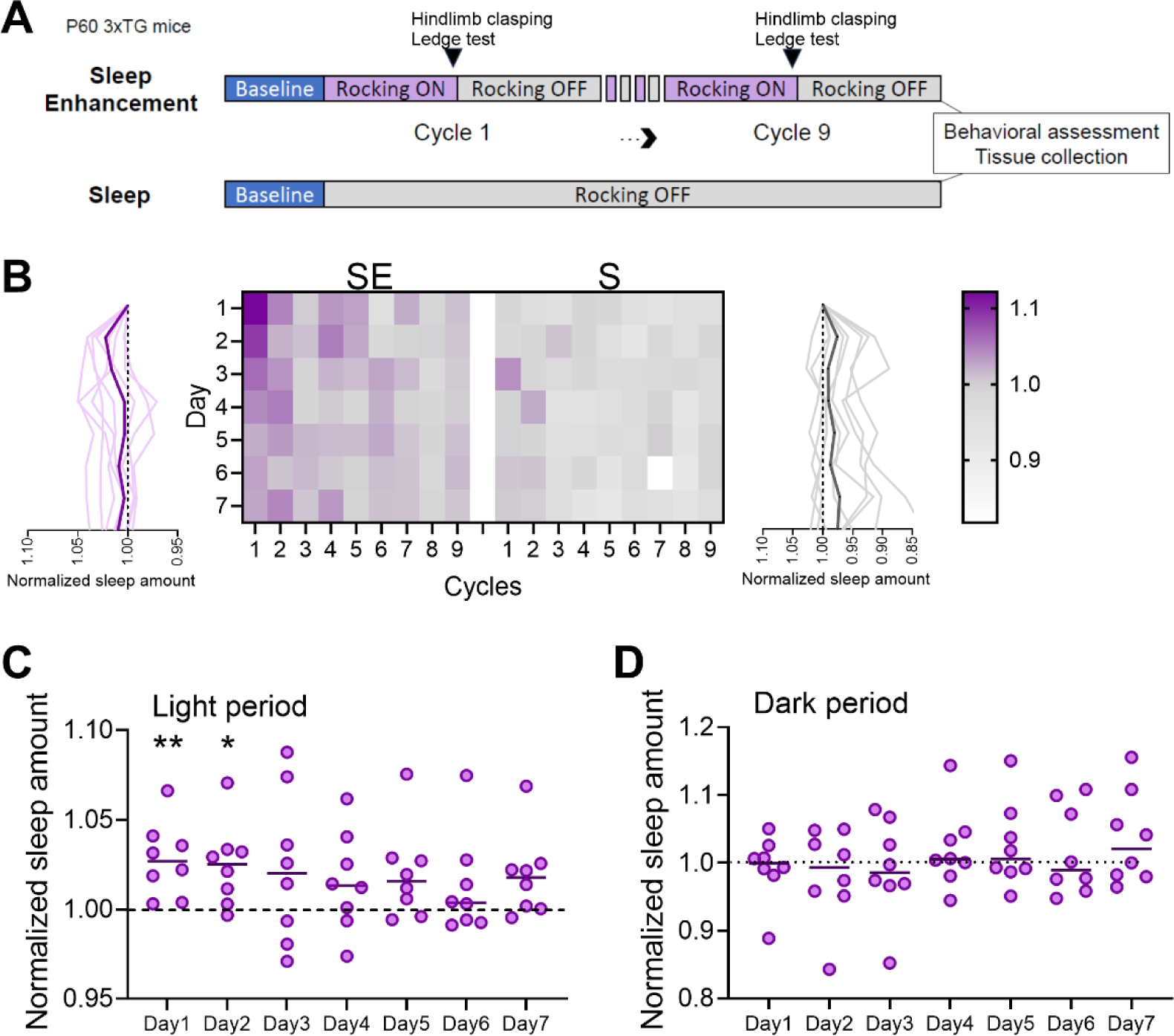
**A**. Experimental design of main study (SE, n=8, S=8). **B**. Heat diagram showing normalized sleep amount for SE and S mice across days and cycles during light period. Graphs on side represent averaged (thick line) and single mouse trends (thin lines) across days. Scale bar shows normalized sleep amount. **C**. Averaged quantification of sleep amount over the 7 days of rocking period (light period only) for SE mice. Values are relative to mean baseline values (dashed line). Each dot represents an individual mouse. *P<0.05; **P<0.01. **D**. Averaged sleep amount over 7 days of rocking period (dark period only) for SE mice.

Moreover, we examined the longitudinal impact of rocking on sleep duration across the nine cycles. Our analysis revealed that the influence of rocking was most pronounced initially and gradually diminished in parallel with aging and disease advancement (RM-1wayANOVA: F (1.732, 10.39) = 7.995 P=0.0094 Figure 4A-B).

**Figure 4.**
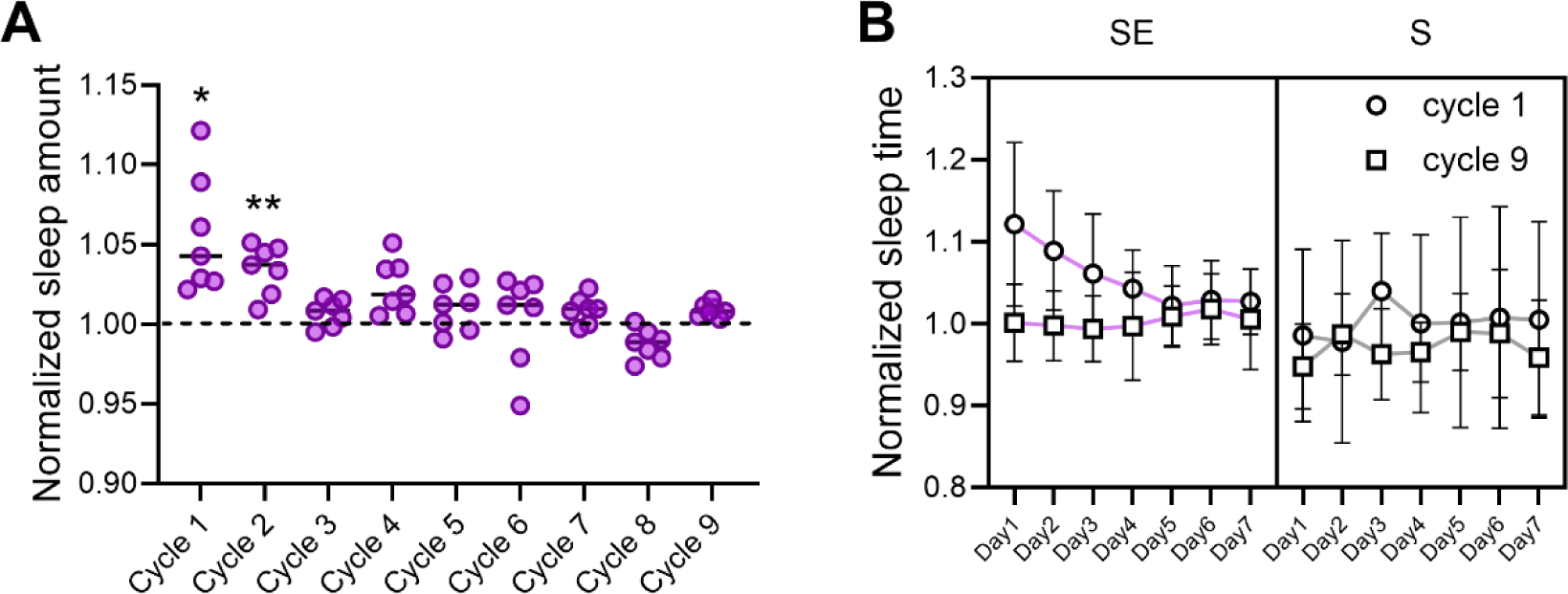
**A**. Averaged quantification of sleep amount over the 7 days of rocking period (light period only) for SE mice. Values are relative to mean baseline values (dashed line). Each dot represents an individual mouse. *P<0.05; **P<0.01. **B**. Sleep amount for cycle 1 and 9 over 7 days of rocking period (light period only) averaged across SE and S mice. Values are mean ± std.

In addition to quantifying sleep amount, we quantified sleep fragmentation by counting sleep-to-wake transitions from the motion activity patterns. This analysis showed that rocking not only prolonged sleep duration but also reduced sleep fragmentation in SE mice (Figure 5A). The pattern of this effect mirrored the trend in sleep duration, being most noticeable at the start of the stimulation week (RM-1wayANOVA: F (2.873, 20.11) = 3.282 P=0.0436) and in the initial cycles (RM-1wayANOVA: F (3.024, 18.15) = 5.155 P=0.0093; Figure 5B-C).

**Figure 5.**
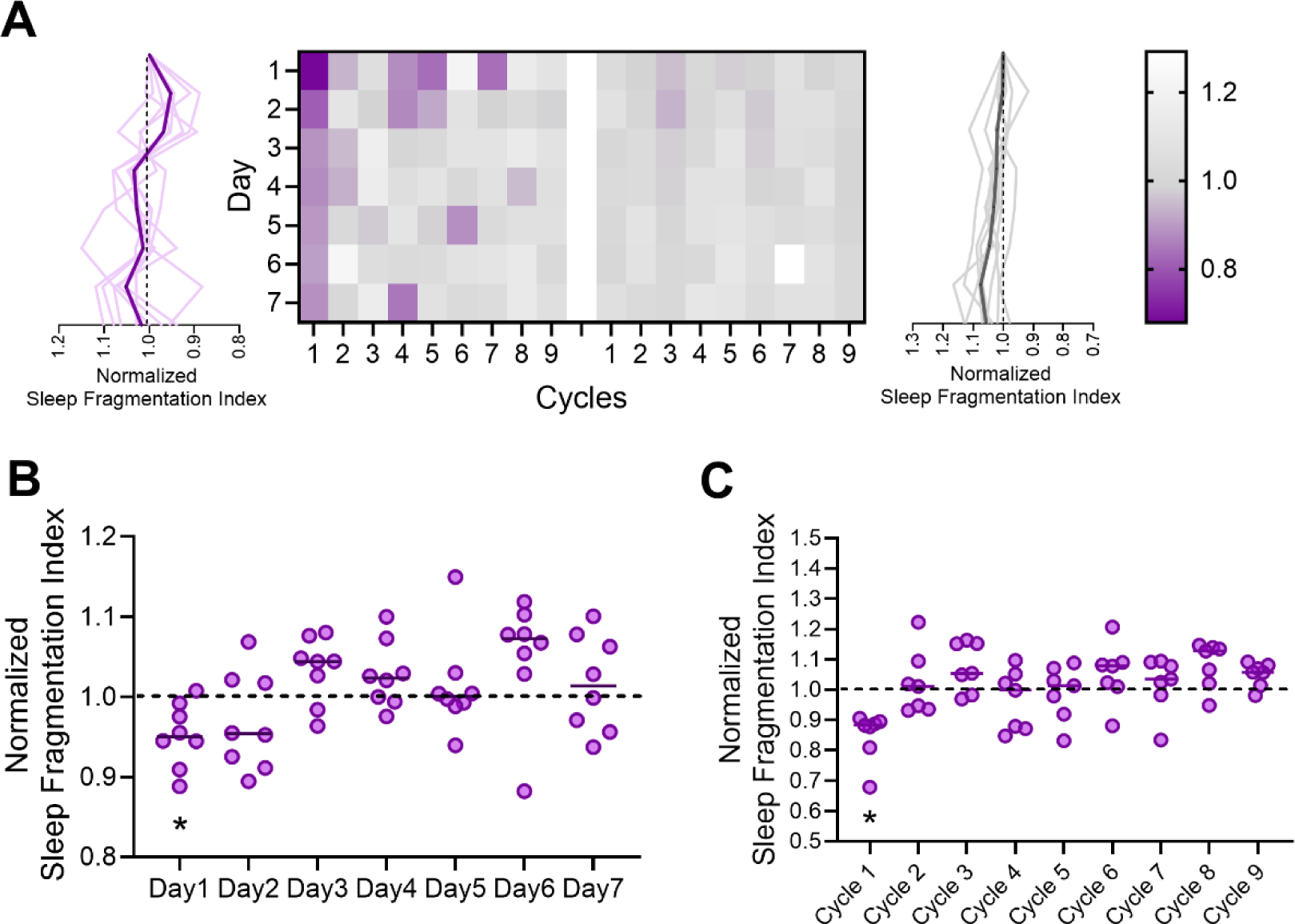
**A**. Heat diagram showing normalized sleep fragmentation index for SE (n=8) and S (n=8) mice across days and cycles during light period. Graphs on side represent averaged (thick line) and single mouse trends (thin lines) across days. Scale bar shows normalized sleep fragmentation index. **B**. Averaged quantification of sleep fragmentation index over the 7 days of rocking period (light period only) for SE mice. Values are relative to mean baseline values (dashed line). Each dot represents an individual mouse. *P<0.05. **C**. Averaged quantification of sleep fragmentation index over the 9 cycle of rocking period (light period only) for SE mice. Values are relative to mean baseline values (dashed line). Each dot represents an individual mouse. *P<0.05.

Thus, rocking enhances both sleep duration and stability. However, this effect is more pronounced initially and diminishes gradually over time, both within individual cycles and across successive cycles of stimulation.

### Rocking delays motor behavior impairment

Throughout the sleep manipulation experiments, mice underwent biweekly behavioral evaluations using hindlimb clasping and ledge tests. These assessments evaluate motor function and coordination and are recognized as indicators of neurodegenerative diseases including AD. Analysis of the scores from these tests over time revealed a decline attributed to disease progression. Notably, scores at the hindlimb clasping test showed no significant effect between S and SE (2wayANOVA: F (1, 210) = 0.2307 P=0.63, Figure 6A), while SE mice outperformed S mice at the ledge walking, a test which requires significant motor coordination (2wayANOVA: F (1, 210) = 9.53 P=0.0023, Figure 6B; unpaired t-test: P=0.004, Figure 6C). The novel object recognition test at the final assessment did not show any significant difference between the groups (unpaired t-test: P=0.5375, Figure 6D). Thus, rocking delayed the progression of motor impairment as evaluated at the ledge walking test but had no effect on learning and memory.

**Figure 6.**
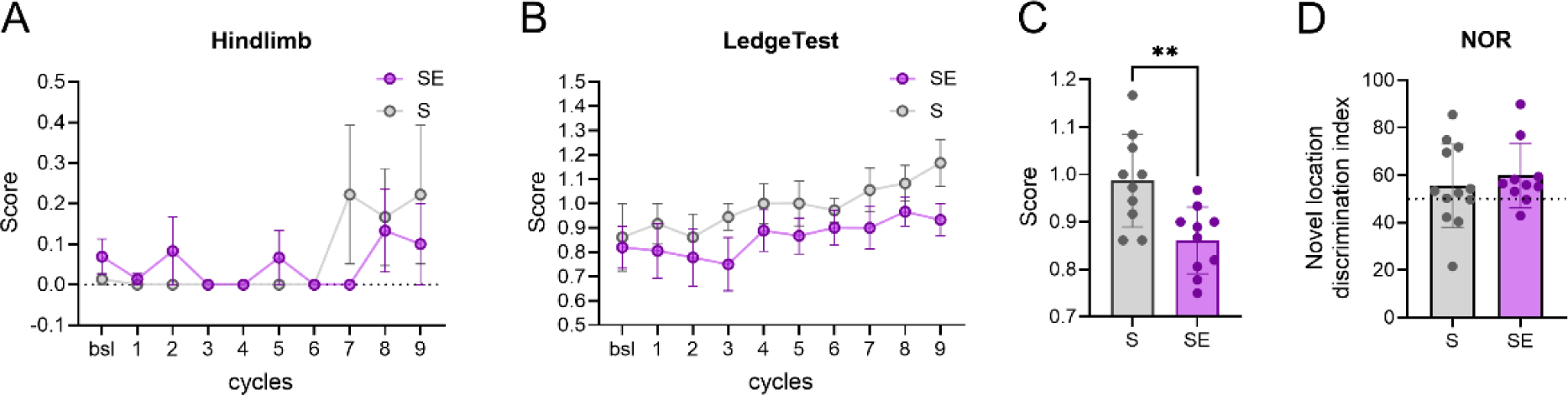
**A**. Averaged scores at the hindlimb test carried out at each cycle (1-> 9). (SE, n=12, S, n=12). Values are mean ± sem. **B**. Averaged score at the ledge walking test carried out at each cycle (1 → 9). (SE, n=12, S, n=12). Values are mean ± sem. **C**. Averaged scores at the ledge walking test across cycles (n=9). **D**. Individual scores of the novel location discrimination index at the novel object recognition test carried out after cycle 9. Dots represent individual mice (SE, n=10, S, n=12), while the dashed line shows when mice spent equal time investigating the old and new objects. In all bar graphs, lines represent mean ± std. **p<0.01

### Rocking reduces beta-amyloid levels

To assess AD pathology, we conducted immunoblot analyses to measure beta-amyloid and tau levels in cerebral cortex samples from SE and S mice. This analysis revealed a significant decrease in beta-amyloid levels in SE compared to S mice (unpaired t-test, P=0.005, Figure 7A). However, examination of beta-amyloid plaques showed no noticeable difference between the two groups, as formations suggestive of plaques were very rare or absent in most of the SE and S mice (Figure 7B). Total tau levels remained largely unchanged between the groups (unpaired t-test, P=0.2474; Figure 7C). In addition, the levels of AT8 and phosphorylated tau at serine 404 showed no significant differences between S and SE (AT8, p=0.26, Figure 7D, p-Tau s404, P=0.58, Figure 7E).

**Figure 7.**
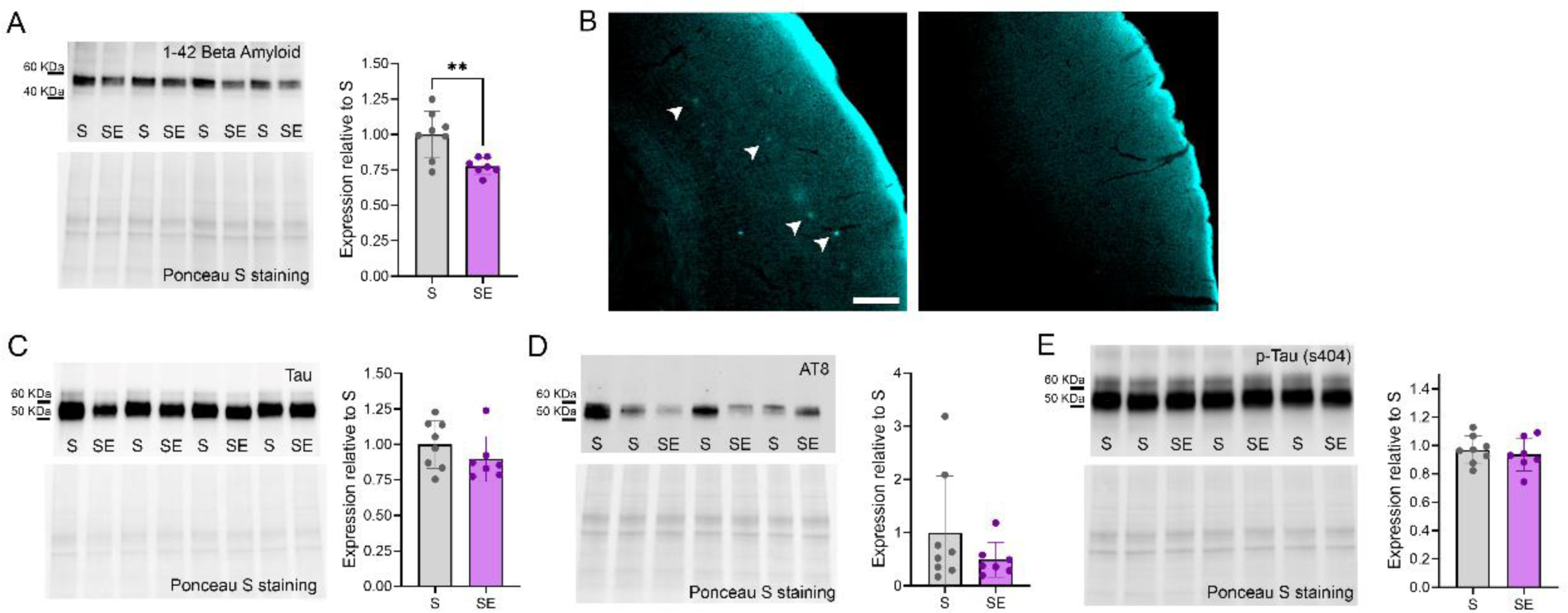
**A**. Representative immunoblot bands stained against beta-amyloid 1-42 and its expression levels in S (n=8) and SE (n=7) mice. **P<0.01. **B**. Representative microscopy fields of cerebral cortex stained with Methoxy-X04 and acquired form a 12-month-old (left) and a S (right) 3xTg mouse. Note that, unlike the aged mouse (left) used a positive control, the S mouse (right) shows no beta-amyloid plaques (arrow heads). Scale bar: 200 µm. **C**. Representative immunoblot bands stained against tau and its expression levels of tau in S (n=8) and SE (n=7) mice. **D**. Representative immunoblot bands stained against AT8 and its expression levels in S (n=8) and SE (n=7) mice. **E**. Representative immunoblot bands stained against p-Tau (ser 404) and its expression levels in S (n=8) and SE (n=7) mice. In all graphs, each dot represents an individual mouse, bars represent mean ± std.

## DISCUSSION

In this study, we demonstrated that rocking 3xTg mice during their sleep period increases NREM sleep duration. However, chronic sleep monitoring revealed that this effect diminishes over time as the mice become habituated to the rocking and as the disease progresses. Despite this, rocking reduced the deterioration of motor behavior over time, which typically occurs in this mouse model. Additionally, chronic rocking during sleep was associated with reduced cerebral cortex levels of beta-amyloid, although it did not significantly affect the expression of total tau and phosphorylated tau.

We found that rocking at 1 Hz during the light period, when the mice normally sleep, led to an increase in time spent in NREM sleep. As observed previously in wild-type animals, this effect occurred at the expense of waking time [19]. Further EEG analysis showed that rocking promoted transitions to short- to medium-duration sleep episodes rather than extending long consolidated sleep, suggesting that, as in humans, rocking facilitates the falling asleep process. The overall increased NREM duration did not change the average amount of SWA, indicating that rocking, unlike other sensory stimulations during sleep (e.g., acoustic stimulation)[26], did not significantly influence sleep intensity. However, when SWA was computed over time, a cumulative effect of rocking was evident due to the greater overall sleep duration. Moreover, rocking did not produce unwanted arousals, even though rocking was not restricted to sleep periods and was always active during the light period.

In this study, we did not utilize a closed-loop process to deliver sensory stimulation exclusively during deep NREM sleep, as is commonly done with acoustic stimulation in human studies [27]. Consequently, we do not know whether applying rocking in such a manner would have produced a stronger effect. However, based on our current data and findings from other studies, it appears that rocking primarily facilitates sleep onset rather than affecting the intensity of preexisting sleep episodes [19,20]. Nonetheless, there is some evidence from human studies indicating that rocking can lead to a faster build-up of delta power, which is indicative of more intense sleep [28].

We found that the effect of rocking on sleep is not constant over time. Even within a single 12-hour period, the sleep-promoting effect of rocking decreases over time, demonstrating a clear habituation phenomenon. This habituation was further observed over multiple days of stimulation. Habituation to sensory stimulation has a long history in research [29]. Early studies by Solokov demonstrated that repetitive patterns of stimulation can induce rapid habituation of the response [30]. This mechanism has been attributed to the reticular formation and is due to a progressive reduction of synaptic efficacy within its circuits [31]. In the case of rocking, habituation could also result from neural adaptation of the vestibular pathways [32]. In our study, we attempted to overcome habituation by implementing a block design, with weeks of stimulation alternated with weeks without stimulation. This approach reduced habituation, as the first days of a new cycle consistently showed better sleep promoting effects than the remaining days. However, the overall sleep promoting effect of rocking declined irreparably over several weeks. It is possible that this decline in effectiveness is not solely due to habituation but also to the progression of the disease and a decrease in overall sleep quality in this mouse model.

Despite the inconsistent effect of rocking on sleep throughout the experiment, we observed some beneficial effects on motor performance. Motor coordination is one of the early deficits in 3xTg mice and is commonly assessed to evaluate disease progression over time [33]. In male mice motor deficit is also more pronounced than in female mice [34,35]. We used the ledge test, which is very sensitive, easy to implement, and can be administered multiple times, to chronically monitor motor performance [23]. Previous studies have shown that chronic sleep disruption can impair motor coordination as assessed by the ledge test in a mouse model of AD [5]. Consistent with these findings, our results indicated that rocking reduced motor performance deterioration, suggesting that motor functions can be better preserved with improved sleep. However, we did not find any effect on the novel object recognition task. This lack of effect may be due to the relatively young age of the animals at testing (P130-140). Cognitive deficits in these mice typically become more apparent around six-eight months of age [36], which could explain the absence of noticeable learning and memory impairment at the time of testing.

In association with improved motor function, we found decreased levels of beta-amyloid in the cerebral cortex of sleep-enhanced mice. This reduction was notable even in the absence of detectable plaques, likely due to the young age of the mice at the time of sacrifice. Numerous studies have linked sleep, or the lack thereof, to beta-amyloid level dynamics and accumulation in both mouse models and humans [8,37–39]. Research has shown that sleep, particularly NREM sleep, facilitates the clearance of beta-amyloid from the brain [9]. During sleep, the glymphatic system is more active, enhancing the removal of beta-amyloid. Conversely, sleep deprivation or disruption has been associated with increased beta-amyloid accumulation [10]. Chronic sleep disturbances can impair the efficiency of the glymphatic system, leading to a buildup of beta-amyloid, which can further disrupt sleep, creating a vicious cycle [10]. Although recent evidence has challenged the role of sleep in clearing the extracellular space from potentially accumulating toxins [40], the evidence supporting sleep in preventing beta-amyloid accumulation remains robust [2]. Consistent with these findings, we observed that sleep enhancement via rocking reduced beta-amyloid levels in the brain. However, we cannot ascertain whether this effect is mediated by improved clearance due to a more efficient glymphatic system or by reduced production of beta-amyloid.

Recent research indicates that chronic sleep restriction significantly increases levels of hyperphosphorylated tau and exacerbates tau pathology progression in both mice and humans [5,6]. Specifically, reductions in slow wave sleep have been associated with tau pathology in individuals with normal cognitive function or mild cognitive impairment [7,8]. However, our findings revealed that sleep enhancement did not affect levels of total tau or phosphorylated tau proteins. One possible explanation for this lack of effect on tau pathology may reside in the animal model used in our study. 3xTg mice typically develop both plaque and tangle pathology, with detectable accumulations of conformationally altered and hyperphosphorylated tau in the hippocampus occurring between 12 to 15 months of age [41]. In contrast, our mice were assessed for tau pathology at a much younger age. It is plausible that the absence of an observable effect of sleep enhancement on tau pathology in our study could be due to the fact that tau pathology had not yet developed at the age when the mice were tested. Another possibility is that modifications in tau protein levels, rather than beta-amyloid deposition, may be more sensitive to changes in sleep intensity. Although direct experimental evidence for this hypothesis is lacking, data from a cohort of human subjects who underwent PET imaging and EEG sleep recordings revealed an inverse relationship between NREM SWA and AD pathology [7]. NREM SWA decreased as evidence of beta-amyloid deposition and tau accumulation increased. Notably, this relationship was stronger with tau pathology than with beta-amyloid pathology [7], suggesting that enhancing SWA, rather than extending NREM sleep duration, may be a more effective approach to positively affect tau pathology.

In conclusion, this study showed that by manipulating sleep via a non-invasive and drug-free methodology based on sensory stimulation it is possible to enhance sleep and modify the trajectory of the disease in a mouse model of AD. These findings also confirm that sleep plays a critical protective role in preventing the progression of neurodegeneration.

## Acknowledgments

We thank Amina Aboufares El Aloui for initial technical support.

## Funding

This work was supported by Alzheimer Research UK (ARUK PPG2020A-023 to MB), Wellcome Trust (215267/Z/19/Z to MB), Armenise-Harvard Foundation (CDA for LdV), ERC-UNICAM (LdV).

## Author contributions

Conceptualization: MB, LdV. Investigation: LZ, LS, NAN, RS, EF, LdV, and MB. Writing—original draft: LZ, LS, LdV, MB. Writing—review and editing: All authors. Project supervision and funding: MB, LdV.

## Competing interests

The authors declare that they have no competing interests.

## Data availability

All data will be made available upon publication.

## Code availability

Codes, accompanied by comprehensive guidelines for replicating the analyses discussed in this paper, can be found on GitHub at https://github.com/BSRLab.

## Notes

### Competing Interest Statement

The authors have declared no competing interest.

### Summary of Updates

Material and Method section has been updated.

